# An Enhancement of Extrachromosomal Circular DNA Enrichment and Amplification to Address the Extremely Low Overlap Between Replicates

**DOI:** 10.1101/2025.06.28.662146

**Authors:** Charles M. Burnham, Alongkorn Kurilung, Visanu Wanchai, Birgritte Regenberg, Jesus Delgado-Calle, Alexei G. Basnakian, Intawat Nookaew

## Abstract

Extrachromosomal circular DNA (eccDNA) of chromosomal origin is present in all eukaryotic organisms and tissues that have been tested. Populations of eccDNA exhibit immense diversity and a characteristically low degree of overlap between samples, suggesting low inheritance of eccDNA between cells or a deficiency in the methods by which eccDNA is detected. This study revisits the Circle-seq approach for enrichment of eccDNA to address these limitations, hypothesizing that experimental procedures significantly contribute to the observed low eccDNA overlap. We optimized the protocol by reducing the time needed to complete the procedure. Linear DNA is digested by increasing Exonuclease V activity. We employed CRISPR-Cas9 for mitochondrial linearization, which proved superior to using restriction enzymes. A key finding is the critical role of random hexamer primer concentration and genomic DNA input in Rolling circle amplification (RCA) for generating high-quality long amplicons from eccDNA (concatemeric tandem copy [CTC]), essential for confident *de novo* eccDNA construction from long-read sequencing data. Lower primer concentrations substantially increased the percentage of CTC-derived eccDNA and improved the overlap of identified eccDNAs in technical replicates. Applying this revised approach to human myeloma and breast cancer cell lines, as well as xenograft models, demonstrated >50% overlap in detected eccDNA, a substantial improvement over the <1% overlap observed in previous studies. Additionally, the oncogenic signature of eccDNAs can be identified across all replicates. These findings provide guidelines for developing standardized procedures for eccDNA profiling, advancing our understanding of eccDNA biology, and its potential clinical applications.

## Introduction

Extrachromosomal circular DNA (eccDNA) of chromosomal origin represents a fascinating and increasingly studied component of eukaryotic genomes (1–4). Distinct from linear chromosomal DNA, eccDNAs exist as separate, covalently closed circles within the cell nucleus that can carry whole or partial genes, regulatory elements, repetitive sequences, and transposable elements. Their specific sequences are highly variable and reflect their genomic origins (1–3). EccDNA gene expression has significant implications for cellular biology, genetics, and disease (1–3). Two defining features of eccDNA populations are their immense diversity and the surprisingly low degree of observed overlap in eccDNA profiles between biological samples and studies (5–12). EccDNA sizes range from a few hundred base pairs (often termed microDNA) to several megabases (1–3) and form through different DNA repair mechanisms (13) in both somatic and germline cells (7,14).

The sequences and the number of specific eccDNA copies vary dramatically from cell to cell, even within the same tissues (5–7,15–18). This multifaceted diversity strongly indicates that the eccDNA complement within a single cell, or a cell population, is a complex and dynamic entity. Consequently, many previous studies have reported a low degree of overlap in eccDNA profiles between different biological and technical replicates of the same tissues (5–9) or cell lines (10–12) based on different eccDNA profiling approaches (9,10). We hypothesize that the current experimental methods are a key factor contributing to the low degree of overlap, due to the high level of heterogeneity among eccDNAs.

Recent advances in enrichment techniques coupled with modern sequencing technologies have been employed to accurately determine eccDNA composition. The dominant enrichment approach, the Circle-Seq method, treats genomic DNA (gDNA) with linear DNA-specific exonucleases to degrade linear chromosomal DNA while enriching circular DNA (19,20). Then, circular DNA of chromosomal origin is further purified by digesting gDNA with restriction enzymes (19,20) or CRISPR-Cas9 (21) to linearize and remove the most abundant form of covalently closed circular DNA in eukaryotic cells, mitochondrial DNAs (mtDNAs), which overshadow eccDNA due to their high copy number. Enriched eccDNA is subsequently amplified using rolling circle amplification (RCA) in preparation for high-throughput short-read or long-read sequencing. Computational tools have been developed to resolve eccDNA structures based on the sequence data through the identification of split and discordant read pairs to identify breakpoints where eccDNAs circularize (22). Notably, with long-read sequencing, the concatemeric tandem copies (CTCs) derived from RCA contain the total context of an eccDNA’s circular conformation and can be captured as a single read, yielding the highest confidence in the construction of eccDNA molecules (20,23,24).

We hypothesize that the current experimental procedure to characterize eccDNA is a key factor contributing to the low degree of overlap in eccDNA profiles. We revised the Circle-seq procedure (19), shortening sample preparation time and significantly increasing the overlap of detected eccDNA across replicates. We also reduced eccDNA enrichment time by increasing the frequency of Exonuclease V supplementation for linear DNA digestion. Furthermore, the CRISPR-Cas9 approach was utilized to linearize mtDNA. We demonstrated that the random hexamer primers concentration in RCA is crucial in generating high-quality CTC reads for the *de novo* construction of eccDNA, and the majority of detected eccDNA was derived from these reads, leading to an increase in overlapping eccDNA. The level of overlapping was further improved by increasing the sampling number. We applied the revised approach to profiling eccDNA from cultured breast cell lines and human myeloma cells and identified highly overlapping eccDNAs that may play a crucial role in cancer biology.

## Materials and Methods

### Cells, cell cultures, and animals

All human breast cell lines were purchased from ATCC. Human cell lines AU565 (RRID: CVCL_1074), HCC1143 (RRID: CVCL_1245), and HCC1395 (RRID: CVCL_1249) were cultured in RPMI-1640 supplemented with 10% FBS. Human SBKR3 cells (RRID: CVCL_0033) were cultured in McCoy’s 5a Modified Medium supplemented with 10% FBS. The MCF10A nontumorigenic human cell line (RRID: CVCL_0598) was propagated and maintained as previously described (28). All cell cultures were cultivated in T75 flasks to 70%-80% confluency and routinely tested for mycoplasma contaminants using a commercially available PCR kit. (Thermo Fisher Scientific). Human OPM-2 (RRID: CVCL_1625) and JJN3 (RRID: CVCL_2078) multiple myeloma cells were provided by G. David D. Roodman, MD, PhD (Indiana University), and Nicola Giuliani, MD, PhD (University of Parma, Italy), respectively, and cultured as previously described (29,30). RPMI 1640 media, fetal bovine serum, Normocin, Plasmocin, and antibiotics (penicillin/streptomycin) were purchased from Invitrogen Life Technologies. For the xenografted animal studies, 7-week-old immunodeficient NSG (RRID: IMSR_JAX:005557) mice were injected intravenously with 10^5^ OPM-2 multiple myeloma cells and sacrificed after 3 weeks, as previously reported (31). Procedures involving animals were performed according to the guidlines of the University of Arkansas for Medical Sciences Institutional Animal Care and Use Committee (protocol #2022200000489).

### Design of guide RNA for mitochondria linearization by CRISPR-Cas9

Single guide RNAs (sgRNAs) were designed via CHOPCHOP v3 (32) under the following criteria: 1) optimal efficiency score (>60), 2) minimal mismatches, 3) no variants within sgRNA target sequence, and 4) all sgRNA shared strandedness to avoid creation of exonuclease-resistant fragments. The target-specific spacer sequences for the sgRNA sequences were as follows: sgRNA1: 5’- ACGGTCGGCGAACATCAGTGGGG-3’, sgRNA2: 5’- TGTTGAGCCGTAGATGCCGTCGG -3’, sgRNA3: 5’- TGAAACCGATATCGCCGATACGG- 3’. All sgRNAs were ordered from Integrative DNA Technologies.

### DNA purification and eccDNA DNA enrichment

We modified the Circle-Seq method as follows: DNA purification was conducted using the Monarch Genomic DNA Purification Kit (New England Biolabs) instead of a plasmid miniprep kit. DNA (1 or 5 µg when specified) was treated with 5 mM NAD^+^, 400 units of Tag DNA Ligase (New England Biolabs), and 1X NEB Buffer to a 50 µL final reaction volume. This reaction was supplemented with 50 units of Exonuclease V (RecBCD) (New England Biolabs), 1 mM ATP, and 1X NEB Buffer 4 to a final reaction volume of 110 μL. Hydrolysis continued at 37°C for 8 hours with the hourly supplementation of 50 units of Exonuclease V, 1 mM of ATP, and 1X NEB Reaction Buffer 4 followed by heat inactivation at 70°C for 30 minutes. Enriched circular DNA was purified by a 1.8x paramagnetic bead (Beckman Coulter) to volume ratio as previously described (33). Aliquots retained at this step served as internal controls.

### Removal of mtDNA via restriction endonuclease

For mtDNA linearization via restriction enzymes, total enriched circular DNA was treated with 30 units of PmeI (New England Biolabs), 1X rCutSmart Buffer, and nuclease-free water, resulting in a final volume of 60 mL. This reaction proceeded for 1 hour at 37°C, followed by heat inactivation at 65°C for 20 minutes. Linearized mtDNA was digested by the addition of two supplements of 20 units of Exonuclease V, 1 mM ATP, and 1X NEB rCutsmart Buffer. Enriched eccDNA was purified by a 1.8x paramagnetic bead to volume ratio as previously described. Experiments comparing linearization techniques were taken from the same pooled DNA enrichment.

### Removal of mtDNA via CRISPR/Cas9 Ribonucleoproteins

Ribonucleoprotein complexes were formed by adding equimolar sgRNA and HiFi Cas9 Nuclease (Integrative DNA Technologies). A reaction of 1 µM sgRNA, 1 µM Cas9 Nuclease, 1X NEB rCutsmart Buffer, and nuclease-free water was allowed to incubate for 20 minutes at 25°C to form ribonucleoprotein complexes, which were then added to the total enriched circular DNA in 1X NEB rCutsmart Buffer for a final 333 nM concentration. This reaction proceeded for 1 hour at 37°C, followed by heat inactivation for 5 minutes at 72°C. Linearized mtDNA was digested as described above. This reaction proceeded at 37°C for 2 hours, with hourly supplementation, followed by heat inactivation for 30 minutes at 70°C. Enriched eccDNA was purified by a 1.8x paramagnetic bead to volume ratio.

### Rolling circle amplification of eccDNA

Enriched eccDNA was amplified by Phi29 polymerase amplification using the phi29-XT RCA Kit (New England Biolabs) according to the manufacturer’s protocol. Random hexameric primers were diluted for RCA optimization, and incubation was extended to 4 hours. Then, eccDNA amplicons (5 µg) were treated with 50 units of T7 endonuclease in 1X NEB Buffer 2 to resolve cruciform structures and incubated for 2 hours at 37°C, followed by 1.8x paramagnetic bead cleaning.

### Acquisition, base calling, data preprocessing, and identification of eccDNA long read data

EccDNA amplicons (2.5 µg) from each sample were prepared for sequencing library creation using the Native Barcoding Kit (Oxford Nanopore Technologies [ONT]) according to the manufacturer’s protocol. Equivalent amounts of barcoded DNA were pooled to ∼1500 ng and used as Ligation Sequencing Kit (ONT) input according to the manufacturer’s protocol. Experiments using DNA purified from breast cancer cell lines were sequenced on R9.4 FLO-MIN106D flow cells on a MinION (Mk1B; ONT). Experiments involving CTC read generation, human breast cancer cell profiling, and myeloma cell lines and xenografts were sequenced on R10.4.1 FLO-MIN106D flow cells using a MinION (Mk1B; ONT). Fast5/POD5 files were demultiplexed and basecalled using Guppy software version 6 or Dorado software version 1.0.0 via the high accuracy model (dna_r9.4.1_450bps_hac_cfg) and (dna_r10.4.1_e8.2_400bps_hac) to generate fastq files. Native barcode adaptors were trimmed using porechop, and reads <200 bp were filtered by NanoFilt (34). The resulting fastq files were processed with CReSIL[16] for eccDNA identification under the following parameters: default settings for the trimming step, relaxed settings for the identification step (-minrsize 50 -break −1), and finally default settings for the identification step.

### Restriction endonuclease recognition site mapping on eccDNA

Consensus fasta eccDNA sequences were extracted from CReSIL output and loaded using the Biopython Library SeqIO module. Sequences were concatenated to confer the circularity, and restriction enzyme recognition sites were searched using the Biopython Restriction module, which can be modified for other restriction endonucleases. Results were compiled in a .txt format, containing the eccDNA ID, the site, and the site position. This output was cross-referenced to CReSIL output based on eccDNA ID, where a positive hit based on ID populated a new column for the restriction endonuclease status. This method was tested on mtDNA, where it correctly identified the PmeI and PacI recognition site.

### Identification of CTC-derived eccDNA

Reads containing eccDNA concatemers were identified in the CReSIL trim step output. Unique CTC read IDs were extracted and used to filter fastq files using seqtk subseq. Resulting fastq files were processed with CReSIL, where CTC-derived eccDNA BED coordinates were compared on a within-sample basis using bedtools intersect querying for IDs that possessed a 90% reciprocal overlap (bedtools intersect -r -f 0.9). These IDs were cross-referenced to the original CReSIL output based on ID.

### Overlapping analysis of identified eccDNAs

All eccDNA genomic coordinates were concatenated into a single master BED file that was subject to comparison, serving as a reference set for all unique eccDNA to be compared. To quantify overlap between each replicate and the master set, we ran bedtools annotate -both optionz, with -i as the master set and -files as individual replicates. Intervals were retained for a given replicate only if all coverage fractions were ≥0.90 (≥90% overlap) These values were used to create a boolean presence/absence matrix indexed by unique interval ID.

## Results

### DNA repair pre-treatment enhances the removal of linearized mtDNA, facilitating the enrichment of circular DNA

Firstly, we reduced the time of linear chromosomal DNA removal from gDNA from 5 days, for the original Circle-seq protocol (19), to approximately 6 hours by adding Exonuclease V cocktail every hour (**Fig. 1A**). Using Exonuclease V, purified from *Escherichia coli* RecBCD, for removal of linear DNA is an important step for eccDNA enrichment. RecBCD is inhibited by modified DNA and can carry out hydrolysis of circular DNA that contains single-strand breaks (25). We then tested whether prior treatment by DNA repair can be used to optimize the removal of linear DNA derived from chromosomes and linearized mitochondria. Using the same DNA pool extracted from HCC1143 cells as a model, we did not observe a difference in the size distribution of unique eccDNAs (**Fig. 1B**) nor the number of eccDNA/Mbp (**Fig. 1C**) due to DNA repair pre-treatment. We observed an increased (1.4%, *P* < 0.05) read utilization for eccDNA construction (**Fig. 1D**), indicating a better digestion of linear DNA in the samples. For mtDNA content, after CRISPR-mediated mtDNA removal, we observed an increased removal (0.4%, *P* < 0.05) of the linearized mtDNA when pretreated with the DNA repair enzyme (**Fig. 1E**). Based on these results, we incorporated a DNA repair step into our eccDNA enrichment to maximize mtDNA removal and enhance eccDNA enrichment. The time required for eccDNA enrichment using our protocol is 15 hours, a significant reduction from the 186 hours for the original Circle-Seq (See Sup. workflow in details).

**Figure 1.**
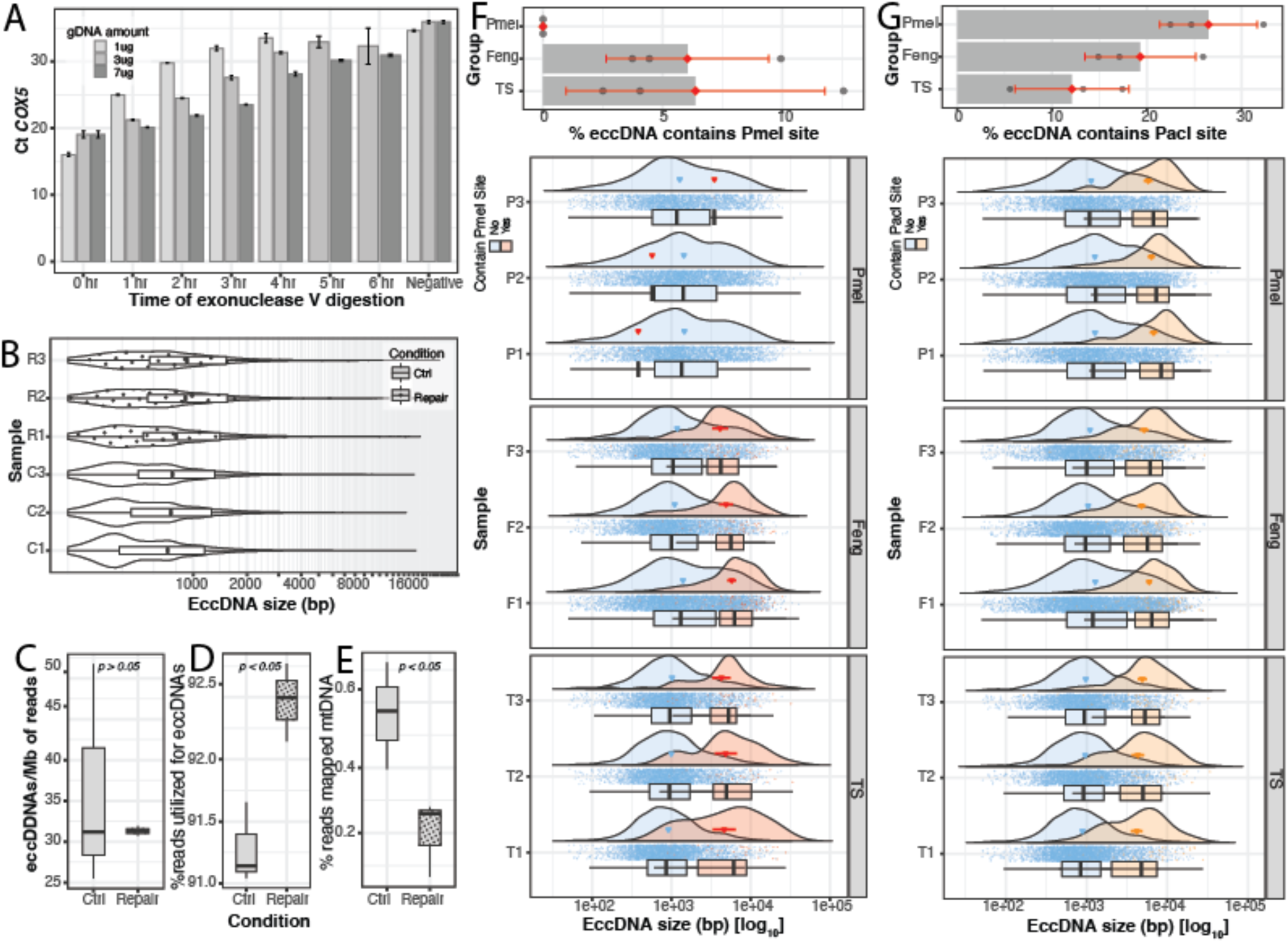
Impact of DNA repair pre-treatment on extrachromosomal circular DNA (eccDNA) enrichment using HCC1143 cells as a model. **A,** Bar plot showing mean raw Ct of COX5 with varying genomic DNA (gDNA) input and time of digestion (*n* = 3). **B,** Violin plot of the size distribution of samples with and without DNA repair. **C,** Bar plot showing mean eccDNA/Mb with and without DNA repair (*n* = 3) **D,** Bar plot showing mean percent read utilization for eccDNA construction with and without DNA repair (*n* = 3). **E,** Bar plot showing the mean percentage of reads mapping to mitochondrial DNA (mtDNA) with and without DNA repair. **F,** Bar plot showing mean percent of eccDNA harboring PmeI cutsite and associated violin plots of eccDNA with (orange) and without (blue) PmeI cutsite (n=3). **G,** Mean percent of eccDNA harboring PacI cutsite and associated violin plots of eccDNA with (orange) and without (blue) PmeI cutsite (n=3). (Student’s t-test where statistics indicated.)

### Targeted mtDNA linearization by CRISPR-Cas9 outperformed restriction enzyme

After chromosomal DNA was removed by Exonuclease V treatment, we aimed to investigate the impact of mtDNA linearization by PmeI restriction enzyme, the most common method, and CRISPR-Cas9 based on sgRNA designed by Feng et al. (21) and our designed sgRNA (see Supplementary Fig. S1 for sgRNA location details), followed by Exonuclease V treatment to remove linearized mtDNA. We tested the removal of mtDNA by qPCR of the mitochondrially encoded Cytochrome C oxidase 1 (*Cox1*), and we observed an over 2-fold qPCR Ct increase in all conditions compared to internal samples before mtDNA removal (Supplementary Fig. S2A), indicating an efficient removal of mtDNA. Sequencing data revealed no significant difference in the percentage of mtDNA reads but a noticeable downward trend when considering the number of cuts to mtDNA with one, two, and three predicted cuts (Supplementary Fig. S2B). The different methods of mtDNA linearization did not result in a difference in the number of unique eccDNAs identified by long read sequencing and mapping (CReSIL) nor in the size distribution of the unique eccDNAs (Supplementary Fig. S2C and S2D). As expected, there were few PmeI recognition sites remaining on eccDNA after treatment with PmeI, but PmeI eccDNA was retained in the CRISPR-mediated conditions (**Fig. 1F** top panel). Interestingly, PmeI eccDNA was larger than non-PmeI eccDNA in the CRISPR-mediated conditions, which is consistent with the probability of the occurrence of a restriction enzyme recognition site increasing when more nucleotides are present (**Fig. 1F** bottom panel). We further assessed the presence of the recognition site of restriction enzyme PacI, which was used for mtDNA removal in previous studies and has more recognition sites in the human genome compared to PmeI (20,24). This experiment resulted in the higher occurrence of PacI recognition sites, approximately twice as much as eccDNA harboring PmeI under CRISPR-mediated conditions (**Fig. 1G** top panel). Also, PacI eccDNA was larger than non-PacI eccDNA (**Fig. 1G** bottom panel).

### Increase CTC read generation by RCA primer titration for high-confidence reconstruction of eccDNA from long read sequences

RCA products containing CTC provide a high confidence of circular amplified DNA molecules. Using the long-read sequencing approach, we can directly identify the CTC characteristic through CTC read, which yields the highest confidence for reconstructing the eccDNA molecule from the sequencing reads. Based on eccDNA profiles from JJN3 cells (**Fig. 1B**), we observed a strong increase in CTC reads that are used to assign eccDNAs with high confidence (CTC eccDNA) (**Fig. 2A**). Under RCA reaction conditions recommended by the manufacturer (50 µM), hyperbranching derived from excess random hexamer primer is a typical strategy to obtain a high yield of amplification product. Therefore, we reasoned that increasing the incidence of CTC reads by reducing hyperbranching during RCA of eccDNA-enriched samples could produce higher-quality reads. We hypothesized that standard reaction conditions from commercially available kits are not optimized for CTC read generation due to the improper ratio of template molecules and substrate random hexameric primers. We then varied the amount of random primer and starting DNA of the same pool of JJN3 cells with two gDNA starting amounts (1 and 5 µg) (recommended by Circle-seq protocol). The amount of RCA product across different amplification conditions correlated with the primer concentration as expected (Supplementary Table S1). Sequencing the products of the varied conditions revealed that lower concentrations of primers consistently increased the percent of CTC reads (**Fig. 2B**) and CTC eccDNA (**Fig. 2C**). A ∼20% increase in CTC eccDNA was observed compared to the widely used standard conditions recommended by the kit. The size of CTC eccDNAs is shorter (**Fig. 2D**), yet their coverage depth is much higher (**Fig. 2E**), reflecting higher abundance, as these molecules were preferably amplified. Consequently, the low input for eccDNA enrichment and lower primer concentrations yielded a higher incidence of eccDNA of common origin and high homology (**Fig. 3F**, Supplementary Fig. 6). With the 90% reciprocal overlap cutoff, the ∼3.5% common origin among three technical replicates of 1 µg gDNA with 2.5 µM primer was substantially improved compared to previous studies reporting extremely low (less than 1%) common eccDNA in both technical and biological replicates (5–11). When considering the correlation of eccDNA to gene density by chromosome, we acknowledged higher correlations with lower concentration (**Fig. 3G** and **H**). Given these observations, we recommend enriching eccDNA from 1 µg of gDNA with 2.5 µM primers for high-abundance eccDNA profiling and enriching eccDNA from 5 µg of gDNA with 5 µM primers for capturing a high eccDNA diversity.

**Figure 2.**
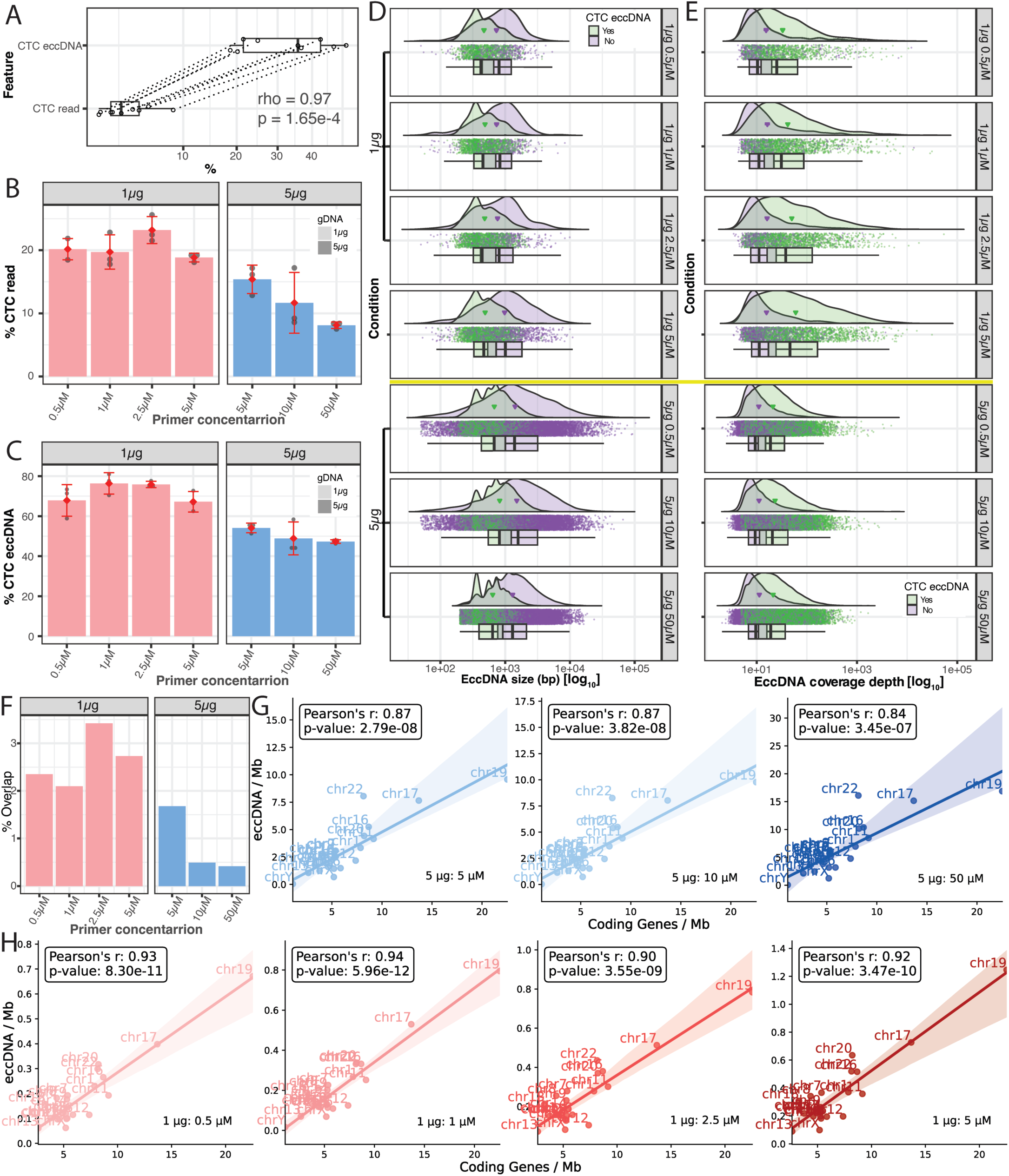
Influence of rolling circle amplification primer titration on extrachromosomal circular DNA (eccDNA) generation using JJN3 cells as a model. **A,** Box and whisker plot showing distribution of and correlation between concatemeric tandem copy (CTC) eccDNA percentage and CTC read percentage (n = 3). **B,** Bar plot showing CTC read percentage at varying primer concentrations using 1 µg input (pink) and 5 µg input (blue) (*n* = 3). **C,** Bar plot showing percent CTC eccDNA at varying primer concentrations under 1 µg input (pink) and 5 µg input (blue) (*n* = 3). **D,** Violin plot showing length of CTC eccDNA (green) and non-CTC eccDNA (purple) using 1 µg input or 5 µg input. **E,** Violin plot showing coverage depth of CTC eccDNA (green) and non-CTC eccDNA (purple) using 1 or 5 µg input. **F,** Bar plot showing percent overlap of replicates at varying primer concentrations using 1 µg input (pink) and 5 µg input (blue) (*n* = 3). **G and H,** Correlation between eccDNA per Mb and Coding Gene per Mb with varying primer concentrations using 5 µg (**G** shades of blue) and 1 µg (**H** shades of pink); points represent the mean between triplicates. Statistical analysis was performed with Pearson’s correlation.

**Figure 3.**
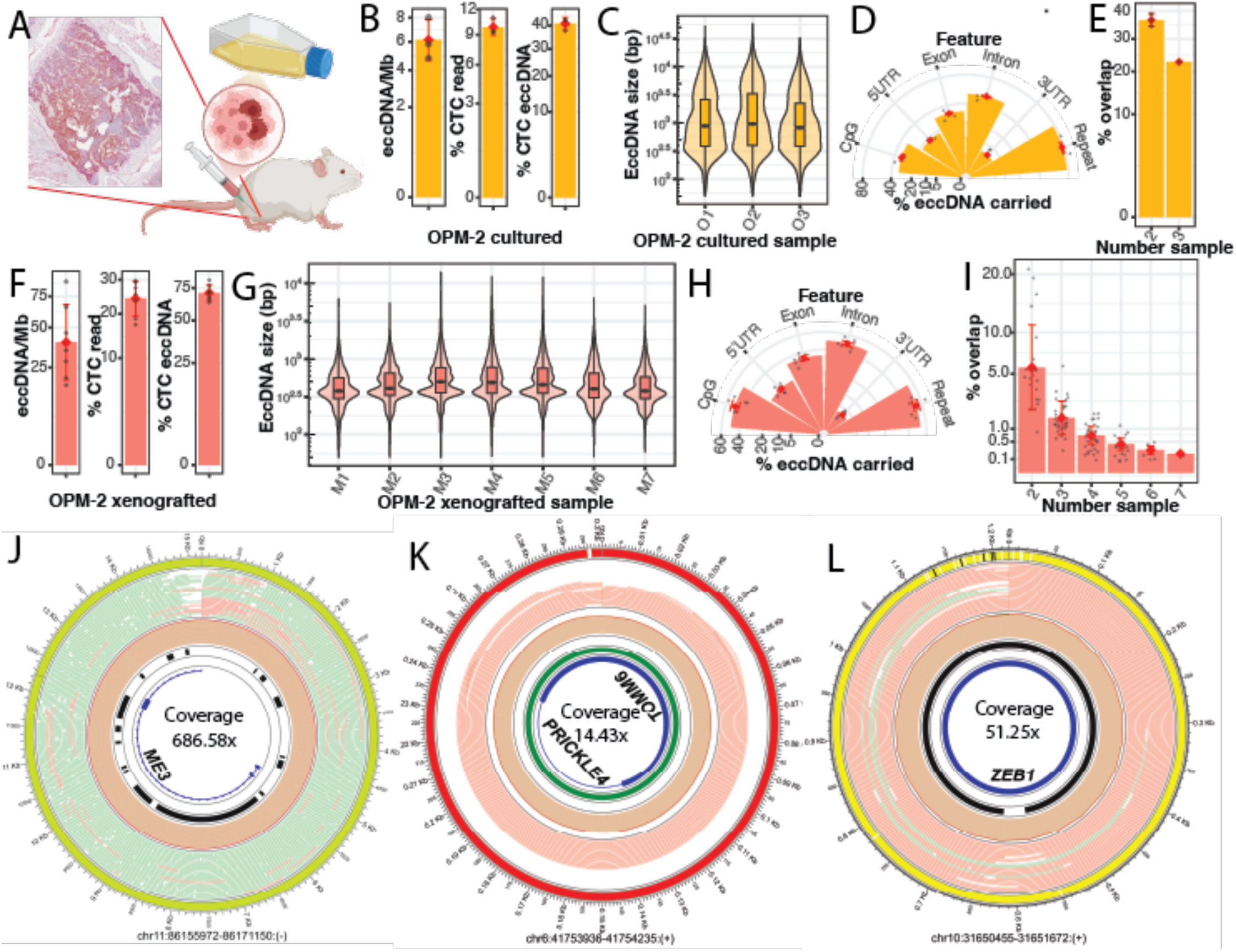
Comparison of extrachromosomal circular DNA (eccDNA) characteristics between OPM-2 myeloma cells cultured and xenografted. **A,** Graphic depicting 7-week-old immunodeficient NSG mice injected intravenously with 10^5^ OPM-2 MM cells. **B,** Bar graphs showing mean eccDNA/Mb, mean concatemeric tandem copy (CTC) read percentage, and mean CTC eccDNA of in vitro cultured OPM-2 cells (*n* = 3). **C,** Violin plot showing the length of eccDNA *in vitro* cultured OPM-2 cells. **D,** Half-moon chart showing percent eccDNA carrying features of in vitro cultured OPM-2 cells (*n* = 3). **E,** Bar plot showing percent overlap of eccNDA between 2 and 3 replicates of in vitro cultured OPM-2 cells. **F,** Bar graphs showing mean eccDNA/Mb, mean CTC read percentage, and mean CTC eccDNA of OPM-2 xenograft (*n* = 7). **G.** Violin plot showing the length of eccDNA in OPM-2 xenografts. **H,** Half-moon chart showing percent eccDNA carrying features of OPM-2 xenografted (*n* = 7). **I,** Bar plot showing percent overlap of eccDNA between of all permutations of OPM-2 xenografted (*n* = 7). **J–L,** Bar plot showing percent overlap of eccDNA between of all permutations of OPM-2 xenografted (*n* = 7). Examples of overlapped eccDNAs identified from cultured OPM-2 cells (**J**), xenografted OPM-2 cells (**K**) and all of them (**L**). See lanes annotation in Supplementary Fig S5.

#### Case study 1: EccDNA profiling of OPM-2 human myeloma cells from culture and mouse xenografts

Next, we applied the enhanced protocol to profile the eccDNA of the OPM-2 human myeloma cell line in DNA samples obtained from the digested tibia of mouse xenografts. With the limited amount of gDNA extracted from mouse bones (∼1 µg), which is a mixture of human cancer cells and mouse cells, we applied 2.5 µM of primers to 1 µg of gDNA to profile the eccDNAs. We obtained detected human eccDNA across seven different samples that widely ranged from 17 to 89 eccDNA/Mbp (**Fig. 3F**), which is significantly higher than the values observed in cell culture experiments. We observed 17%–30% of CTC reads, which resulted in 63%–80% CTC eccDNA (**Fig. 3F**) across different samples. The identified eccDNA distribution across samples was similar (**Fig. 3G**) and shorter than that in the cell culture experiments (**Fig. 3C**), possibly due to the starting gDNA amount. We observed a high correlation of eccDNA to gene density by chromosome. (Supplementary Fig S3). We observed across the seven samples that, on average, the identified eccDNAs carried 50% CpG, 20.7% 5’ UTR, 34.4% exon, 49.5% intron, 3.7% 3’ UTR, and 52.9% repeats (Fig. 3H). The percent overlap of human eccDNAs in xenografted OPM-2 cells (**Fig. 3I**) was significantly lower compared to cultured OPM-2 cells (**Fig. 3E**), with an average 6.5% pairwise overlap and a 0.19% seven-sample overlap. We arbitrarily selected overlapped eccDNAs found in the three cultured OPM-2 cell biological replicates, the seven biological replicates of xenografted OPM-2 cells, and the ten samples combined. One of the selected, commonly found eccDNAs in OPM-2 cultures was a >15 Kbp non-CTC eccDNA (**Fig. 3J**) with a >680x coverage depth, originating from chromosome 11, carrying three exons of the *ME3* gene and many repeat elements. Another selected eccDNA was a small (∼300 bp) CTC eccDNA (**Fig. 3K**) with a 14.43x coverage depth, originating from chromosome 6, that was found in common among seven replicates of xenografted OPM-2 samples. We found that this eccDNA contained an exon of the TOMM6 gene with an intron from the *PRICKLE4* gene. Notably, this eccDNA was entirely derived from CpG sequences. Interestingly, we found some common eccDNAs among all the samples (**Fig. 3L**). One common CTC eccDNA had a size >1.2 Kb with a 50x coverage depth, originating from chromosome 10 and carrying an exon of the transcription factor *ZEB1* gene.

#### Case study 2: EccDNA profiling in human breast cells

We first performed eccDNA profiles of HCC1143, HCC1395, AU565, and SKBR3 cultured human breast cancer cells using the same procedure described for profiling cultured OPM-2 cells (5 µg of gDNA with 5 µM primers). We found a very low overlap (<5%) among biological replicates of all the cancer cells (**Fig. 4A**), unlike cultured OPM-2 cells (>20% overlap; **Fig. 3A**). To approach the high heterogeneity of eccDNAs reported in many previous studies, we further increased the number of samples by processing two technical replicates of the same gDNA aliquot then combined them for profiling of eccDNA in the context of biological replicates. We applied 2.5 µM of primers to 1 µg of human breast cancer cell lines gDNA, as well as to MCF10A epithelial cells cultured under similar conditions. Using this strategy, we obtained the average of eccDNA/Mbp with replicates varying from 3.8 to 8.2 eccDNA/Mbp (**Fig. 4B**). We observed 5%–20% of CTC reads (**Fig. 4C**), which generated 38%–66% CTC eccDNA (**Fig. 4D**) across different cell lines. The distribution of the identified eccDNA among the cell lines was slightly different (**Fig. 4E**). As expected, we observed a high correlation between eccDNA and gene density by chromosome. (Supp. Fig. S4). Assessing the genomic content of eccDNA may provide evidence of its biogenesis. With similar genomic content across the cells, a significant proportion of eccDNA was enriched for CpG, 5’UTR, exon, and intron, indicating that the biogenesis of eccDNA likely favors regions of high transcriptional activity (**Fig. 4F**). As previously reported, eccDNA is enriched by repeat elements, which favors the formation of eccDNA through microhomology (**Fig. 4F**). In analyzing the breast cell models for eccDNA of both common genomic origin and homology, we first assessed the eccDNA’s overlap between two technical replicates over different biological replicates. We observed as high as 18.9% (SKBR3 biological replicate 2)–50.4% (HCC1143 biological replicate 1) of overlapped eccDNAs (**Fig. 4G**). The overlap of eccDNAs among the three biological replicates of individual cells was quite high—19.5% for SKBR3, 30.1% for HC1395, 31.8% for AU565, 42.2% for MCF10A, and 43.4% for HC1143 (**Fig. 4H**). We arbitrarily selected an example of overlapped eccDNA found among the three biological replicates of individual cultured MCF10A breast cells and identified a CTC eccDNA (>1.5 Kbp) (**Fig. 4I**) with a 70x coverage depth originating from chromosome 2. This eccDNA contains a *BC16143* gene exon of and an *LRP4* gene intron with a long repeat. Another commonly identified non-CTC eccDNA among replicates of AU565 cells was >9.5 kbp, with a high >920x coverage depth, originating from chromosome 16. We found that this eccDNA contains an *MYLK3* gene exon with numerous repeat elements (**Fig. 4J**). We also found a higher coverage depth (>1900x) of CTC eccDNA as a common feature among replicates of HCC1143 cells (**Fig. 4K**). This eccDNA was >1.3 Kbp, derived from chromosome 10, and carried an intron-containing microRNA 2 repeats of the *ENTPD1* gene. An overlapped eccDNA replicate of HCC1143 cells derived from chromosome 5 that was >6Kbp with an 868x coverage depth (**Fig. 4L**) contained 5 repeats, a CpG, and part of the *TCF7* gene. Lastly, we found an over 9.5Kbp non-CTC eccDNA (519x coverage depth), originating from chromosome 15, carrying many exons and introns of the *POLG* gene and many repeats commonly identified among SKBR3 replicates (**Fig. 4M**).

**Figure 4.**
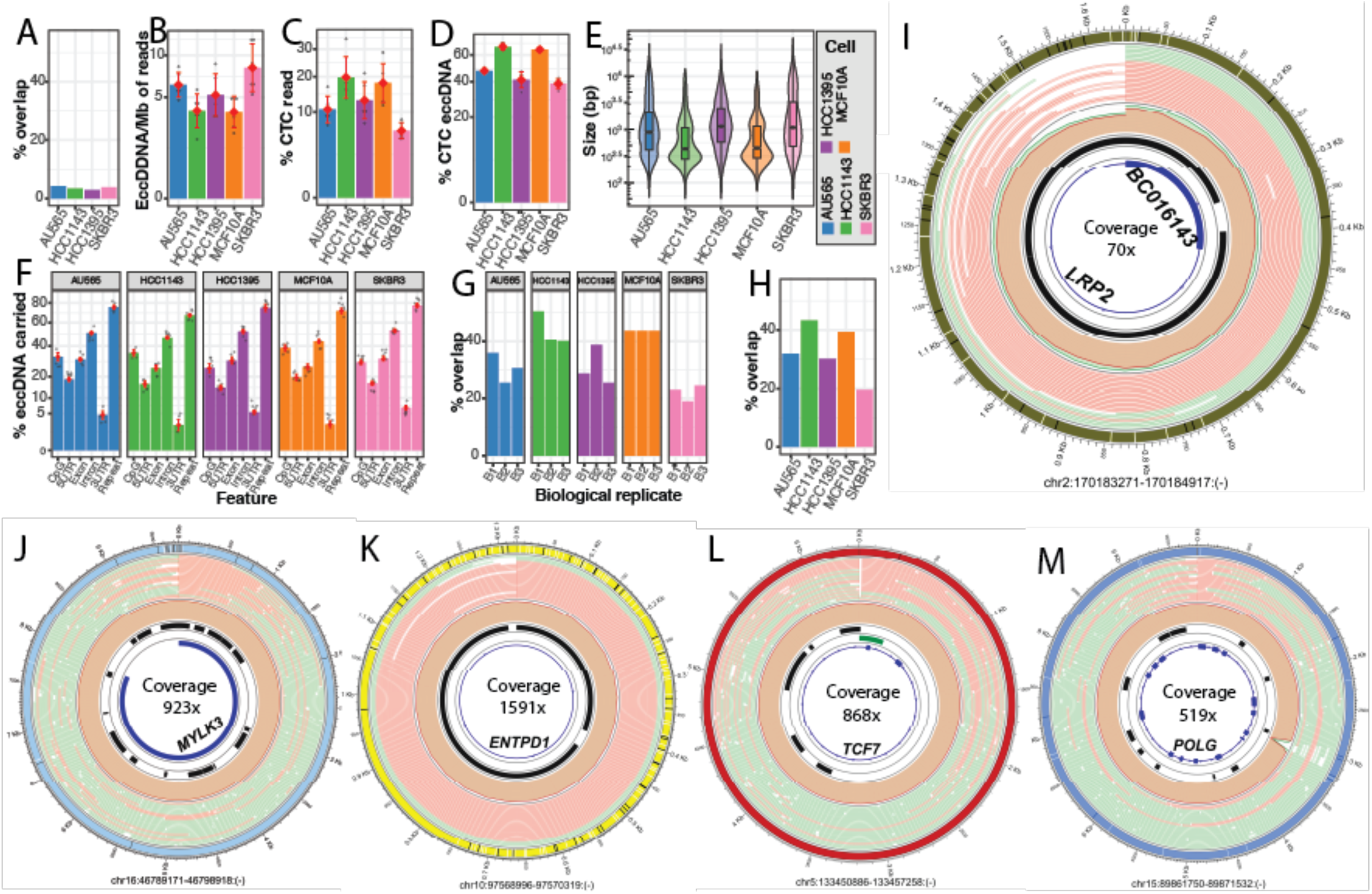
Profiles of extrachromosomal circular DNA (eccDNA) across different human breast cancer cell lines. **A,** Bar graph showing percent overlap between replicates of breast cancer cell lines. **B,** Bar plot showing mean eccDNA/Mb of breast cancer cell lines (*n* = 3). **C,** Bar plot showing mean mean concatemeric tandem copy (CTC) read percentage of breast cancer cell lines (*n* = 3). **D,** Bar plot showing mean CTC eccDNA percentage of breast cancer cell lines (*n* = 3) **E,** Violin plot showing average length of eccDNA of breast cancer cell lines (*n* = 3). **F,** Bar plot showing mean percentage of eccDNA carrying selected features in breast cancer cell lines (*n* = 3). **G,** Bar plot showing mean overlap between combinations of technical replicates of breast cancer cell lines. **H,** Bar plot showing overlap between 3 biological replicates of breast cancer cell lines. **I–M,** Examples of overlapped eccDNAs identified from MCF10A cells (**I**), AU565 cells (**J**), HCC1143 cells (**K**), HCC1395 cells (**L**) and SKBR3 cells (**M**). See lanes annotation in Sup. Fig S5.

### Oncogenic signatures of eccDNAs identified in multiple myeloma and breast cancer

We further investigated the curated 758 oncogenes list from the COSMIC database (version 102) (26) to examine whether their components were carried by the identified eccDNAs derived from the samples of the two case studies. For xenografted OPM-2 cells, we observed 8 common oncogenes from the detected 358 oncogenes, distributed in 1557 identified eccDNAs across the 7 replicates (**Fig. 5A**; see supplementary data for details). For cultured OPM-2 cells, we observed 52 common oncogenes from the detected 182 oncogenes, distributed in 1557 identified eccDNAs across the 3 replicates. The breast cells had many fewer detected oncogenes than the multiple myeloma cells. In the breast cell family, SKBR3 cells had the most detected oncogenes (143 genes) distributed in 326 identified eccDNAs across 3 replicates, with 21 oncogenes common among the 3 replicates. The nontumorigenic MCF10A epithelial cells had the fewest detected oncogenes, 17 distributed in 60 identified eccDNAs, with 7 in common among the 3 replicates. Subsequently, the basic characteristic features of eccDNAs carrying oncogenes are illustrated in **Fig. 5B** (see supplementary data for details). A strong association is clearly seen between the commonly detected oncogenes across replicates, the number of eccDNAs carrying those oncogenes, and the coverage depth in all cells. Interestingly, we found some common oncogenes between xenografted and cultured OPM-2 cells (highlighted in a thick border in **Fig. 2B**). *NOTCH1,* encoding notch receptor 1, which is a well-known oncogene (27) in multiple myeloma, was detected in all seven replicates of xenografted OPM-2 cells, where it was carried by 40 eccDNAs with a maximum coverage depth of 180x. It was also detected in all three replicates of cultured OPM-2 cells, carried by seven eccDNAs with an extremely high maximum coverage depth of over 88,000x. *PRDM16*, which encodes a transcription factor that plays a crucial role in various cellular processes, was detected in all 7 replicates of xenografted OPM-2 cells, carried by 65 eccDNAs with a very high maximum coverage depth >2,100x. It was also detected in all 3 replicates of cultured OPM-2 cells, carried by 17 eccDNAs with a high maximum coverage depth >668x. *PRDM16* was also a top-ranked oncogene in SKBR3 cells (bottom right panel) and was carried by 13 eccDNAs with a high maximum coverage depth of >864x. Moreover, we also found that *PRDM16* was detected in the rest of the breast cells, appearing in two AU565, two HCC1143, three HCC1395, and two MCF10A replicates.

**Figure 5.**
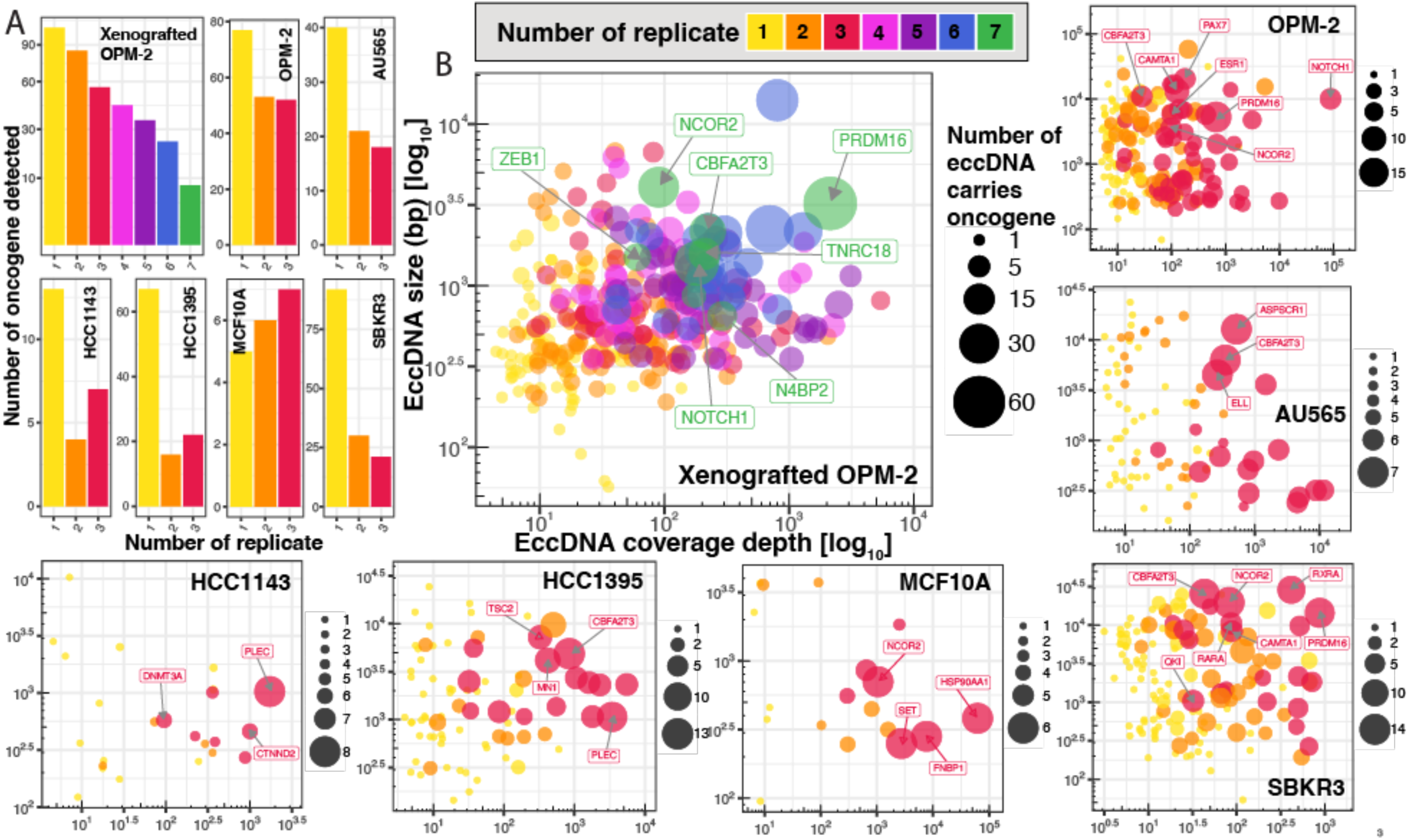
Oncogene distribution analysis of identified extrachromosomal circular DNA (eccDNA) from the two case studies (Figs. 3 and 4). **A,** Bar plots show the number of oncogenes, or their components detected on eccDNAs identified across several replicates of different cells. **B,** Bubble plots showing the basic characteristics (size on the y-axis and coverage depth on the x-axis) of eccDNAs carrying an oncogene or its components. The dot size represents the number of eccDNAs that carry a specific oncogene across different replicates. Name tags with a gray arrow indicate top-ranked oncogenes (large dots) found in all replicates. The figure annotations are the same for all individual plots. In all plots, the color indicates the number of replicates in which an oncogene was detected on the identified eccDNAs.

## Discussion

Our revised protocol demonstrates that eccDNA enrichment can be conducted in as little as 8 hours, representing a significant reduction compared to traditional protocols that require up to 7 days (19). This enhanced method can potentially benefit clinical applications that require timely results. It has been suggested that mtDNA removal is necessary to capture longer eccDNA in colorectal tumor samples (21). Studies indicate that without mtDNA removal, upwards of half the reads represent mtDNA (12). Traditional procedures utilize restriction enzymes to linearize mtDNA as part of the eccDNA enrichment process (19,20), indicating that the removal of mtDNA is a crucial step for eccDNA enrichment.

Our study determined that longer eccDNAs are more likely to be lost when restriction enzymes are used to remove mtDNA. Although termed rare-cutting in the context of other restriction enzymes, these enzymes still possess a high probability of cutting human chromosomal DNA. Given equal nucleotide content, an 8-mer restriction enzyme is predicted to cut roughly every 65 kilobases. With this probability, one can expect that a longer eccDNA is more likely to contain a recognition site for the restriction enzyme. This trend was observed among eccDNA that includes the recognition site for both PmeI and PacI. Our findings were contrary to a recent study comparing eccDNA enrichment techniques, in which the authors suggested that eccDNA harboring a recognition site was smaller (12). This conclusion was not supported by our findings, which showed that eccDNA containing either a PmeI or PacI recognition site had a longer size profile than those that did not contain a recognition site. Given the length of sgRNA used for mtDNA linearization, it is very unlikely that larger eccDNA will contain a target sequence for such an sgRNA. Our results highlight the need to consider the selection of mtDNA linearization techniques. Identifying genes related to genomic stability in breast cancer, such as MTOR, AKT1, and SMAD3, harboring eccDNA, demonstrates the need to screen for genes of interest before selecting an mtDNA linearization technique (See Supplementary Table S2).

RCA for eccDNA amplification is the preferred approach, as it creates split and discordant reads used to assemble eccDNA by identifying breakpoints, and protocols and software suites have been developed to exploit this and were recently compared (10). Due to the high processivity of RCA, eccDNA amplicons can span the circumference of the circle multiple times, resulting in CTC reads. As part of the library preparation for short-read sequencing, these CTC reads are fragmented to meet the read length requirement. In long-read sequencing, the length of the DNA fragment is characteristic of the read length. Therefore, amplicons from RCA are represented in nanopore sequencing data and can be used for the high-confidence identification of eccDNAs. Previous studies did not seek to optimize RCA for the generation of these reads, as they could not be detected using short-read sequencing technology. We found that the manufacturer’s recommended random-hexameric primer concentration is suboptimal for generating CTC reads. Given our workflow, a lower random-hexameric primer concentration, paired with lower gDNA input, consistently increased CTC read incidence. We recommend that researchers perform pilot experiments to identify the specific conditions of primer concentrations and input, depending on their availability, as we found that different cell lines and conditions have different levels of eccDNA characteristics and heterogeneity, as previously reported (16).

Low amounts of starting gDNA and primers in RCA can result in over half of the eccDNA being derived from CTC reads, as these reads participate in the *de novo* assembly of the majority of eccDNA with high coverage depth. However, long eccDNAs will be rare compared to the high amount of starting gDNA. Therefore, the amounts of gDNA and primer of choice for RCA will depend on the study’s objective—capturing a wide range of eccDNA heterogeneity or capturing a high-abundance of eccDNAs. Nevertheless, our findings revealed a high overlap of identified eccDNAs, which is obtainable under proper reaction conditions, unlike previous studies that reported less than 1% overlap (5–12). Within the high degree of eccDNA overlap achieved by the enhanced procedure, we also found common oncogenes detected across replicates. These common signature oncogenes were carried by many eccDNAs with high coverage depth. This supports the known, strong link between eccDNA and oncogene amplification and overexpression in cancer. (1).

The phenomenon of “low overlap” in eccDNA research poses a complex challenge that significantly hinders the consistent detection, characterization, and functional understanding of these diverse genetic elements, as well as the development of biomarkers for clinical diagnostics and therapeutics. This study demonstrated the technical importance of steps in eccDNA enrichment and its application to increase the overlap level of the identified eccDNAs based on long-read sequencing technology. The results from this study provide the basis for the development of a community-accepted procedure for eccDNA profiling in the future, advancing our understanding of eccDNA biology and its implications.

## Data availability

All sequencing data from 79 samples were deposited and made publicly available in the NCBI SRA database under BioProject PRJNA1283158. The update of the CRSIL software (version 1.2.0) to be compatible with the R10 chemistry of Oxford Nanopore sequencing and an additional report on CTC-related features are available at https://github.com/visanuwan/cresil.

## Funding

Musculoskeletal (MSK) Health and Disease Creativity Hub of the UAMS College of Medicine, P20GM125503, T32GM106999, R37CA251763

## Notes

### Competing Interest Statement

The authors have declared no competing interest.

### Summary of Updates

The manuscript has been proof read and edited to improve the language.

