## Supplementary for "An Enhancement of Extrachromosomal Circular DNA Enrichment and Amplification to Address the Extremely Low Overlap Between Replicates"

**Figure S1**

Circos plot of mitochondrial genome. Tracks from outside in: 1) Tile representing mitochondrial genome with bands relative to features in center, 2) Gene annotation (white blocks) with SNPs from variant calling (red dashes), 3) Line plot representing 50 base pair rolling depth from native sequencing, 5) Feature relative to tiles in track 1.

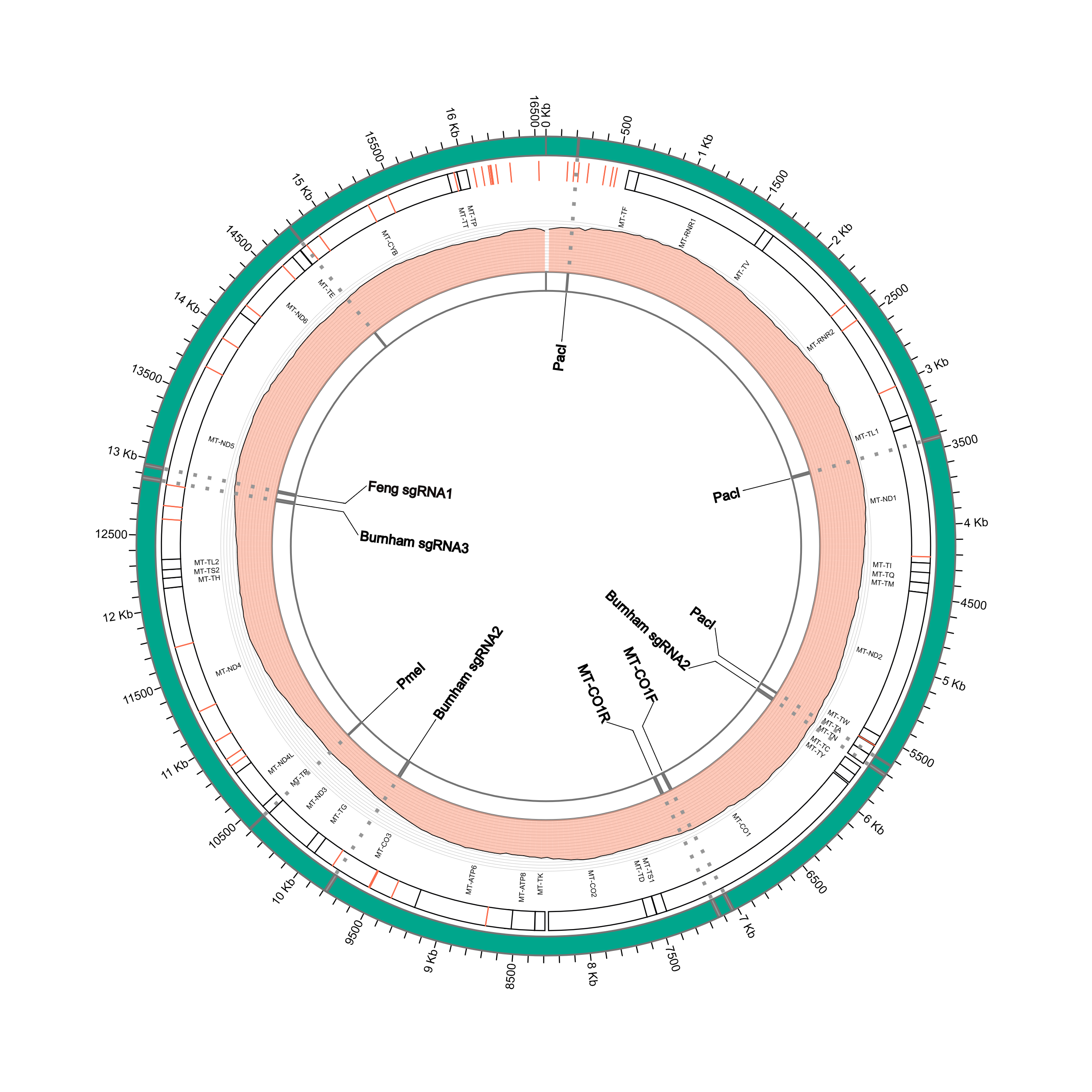

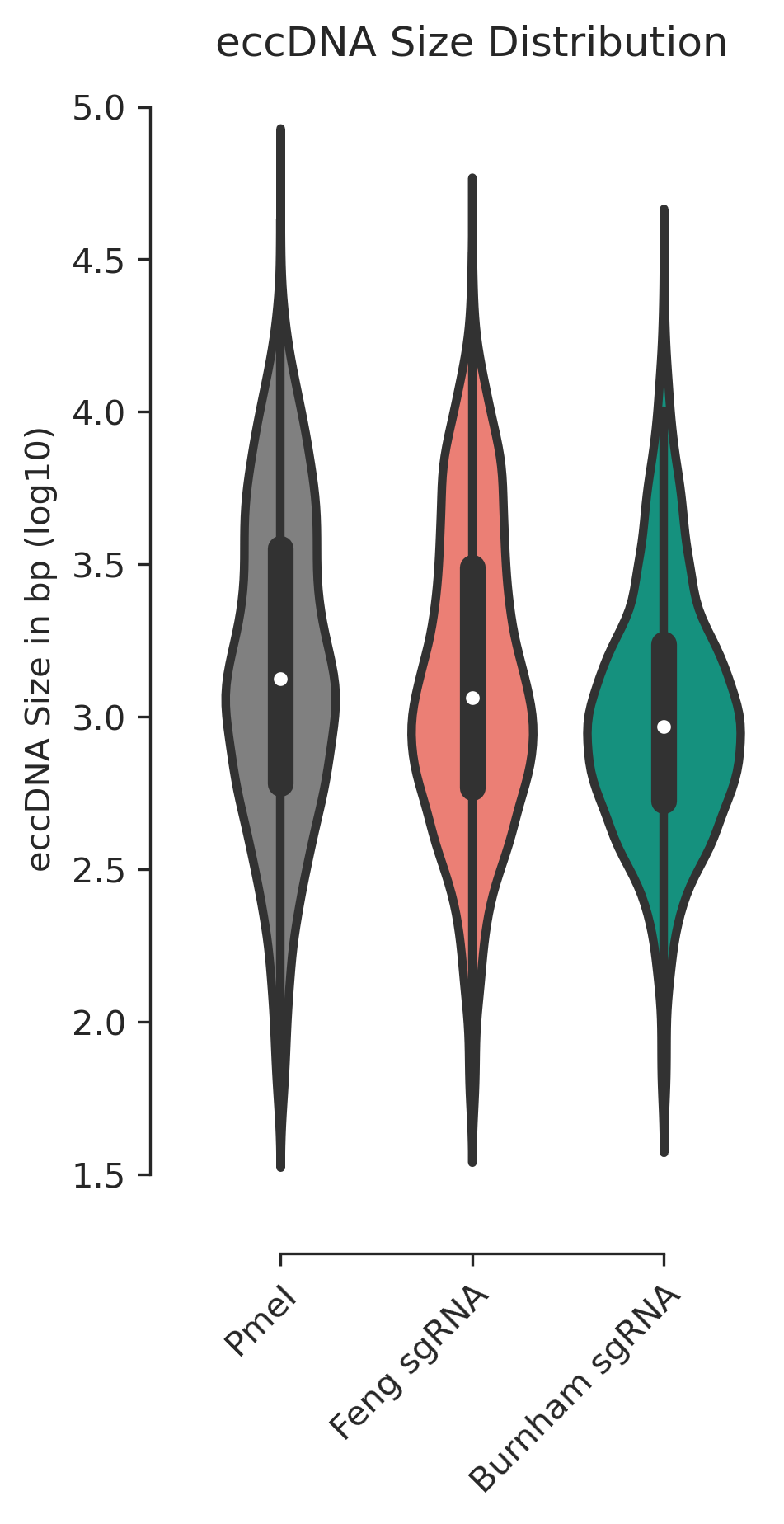

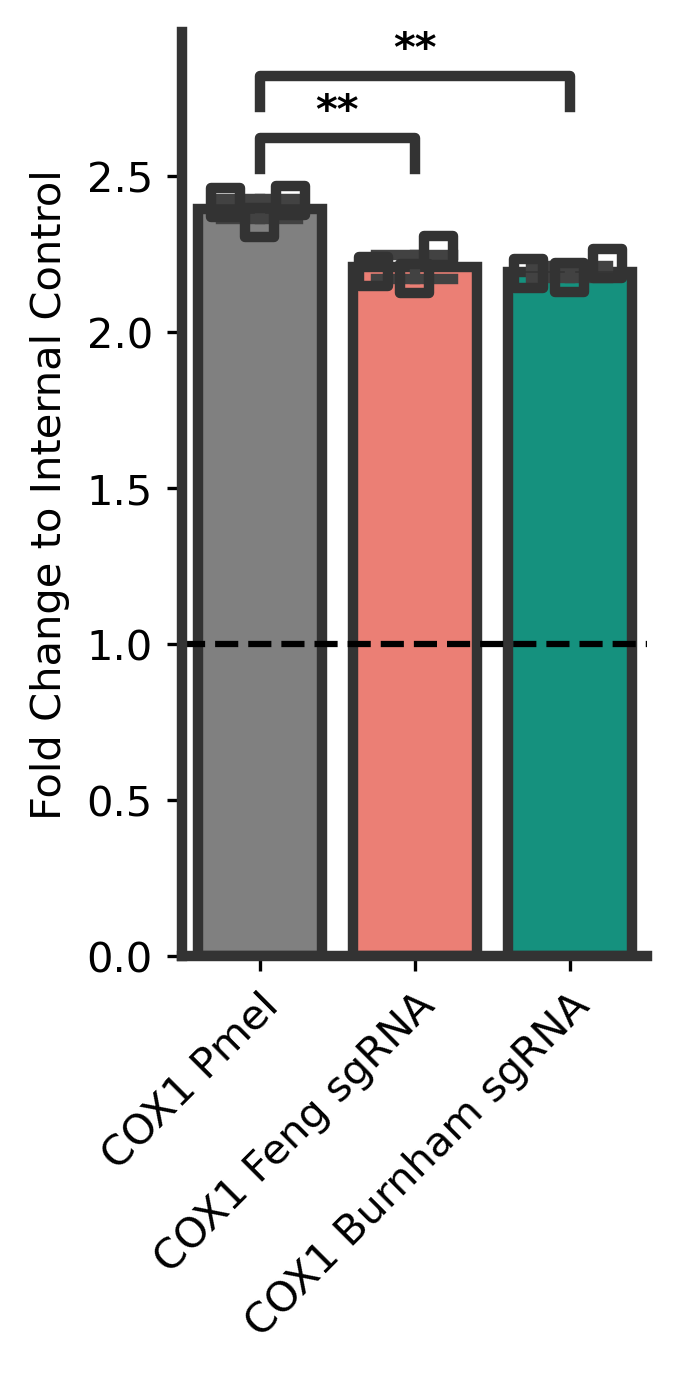

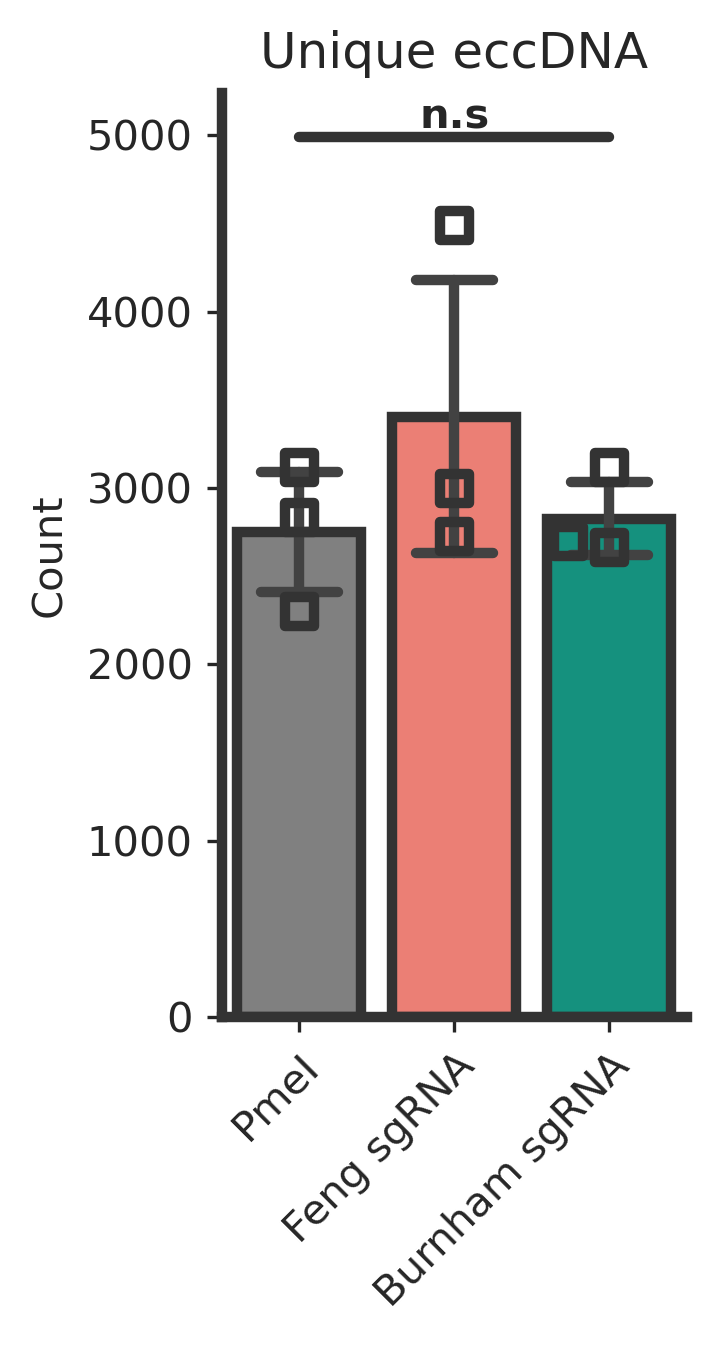

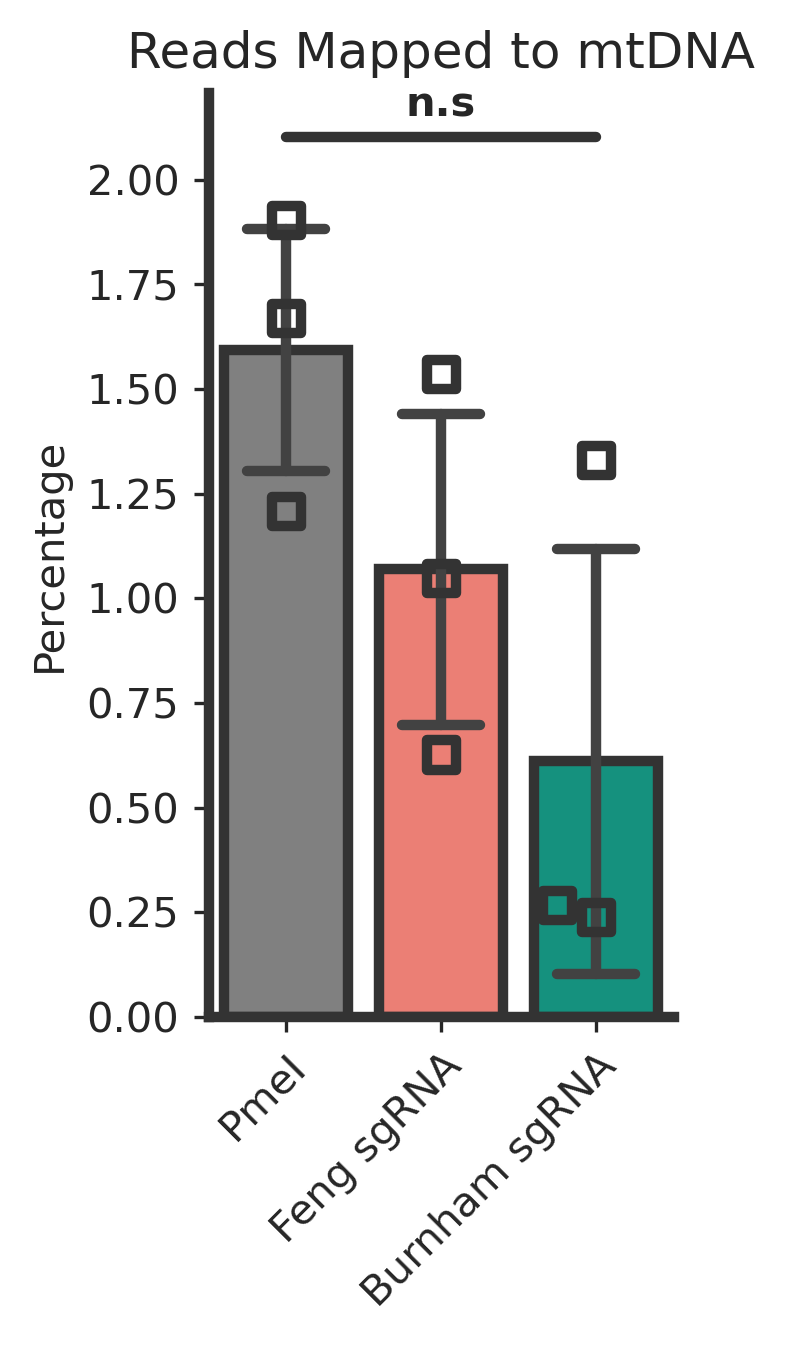

**Figure S2**

**CRISPR-mediated mtDNA linearization. A,** Mean fold change of COX1 comparing mitochondrial DNA (mtDNA) linearization technique (n = 3). **B,** Mean percent reads mapped to mtDNA by condition (n = 3). **C,** Mean count of unique eccDNA detected by CReSIL per condition (n = 3). **D,** Violin-box distribution of identified eccDNA. Asterisks indicate significant differences (**P* < 0.05; ***P* < 0.01; *****P* < 0.0001; ns: not significant, ANOVA for group comparison, Tukey’s HSD test for pairwise comparison)

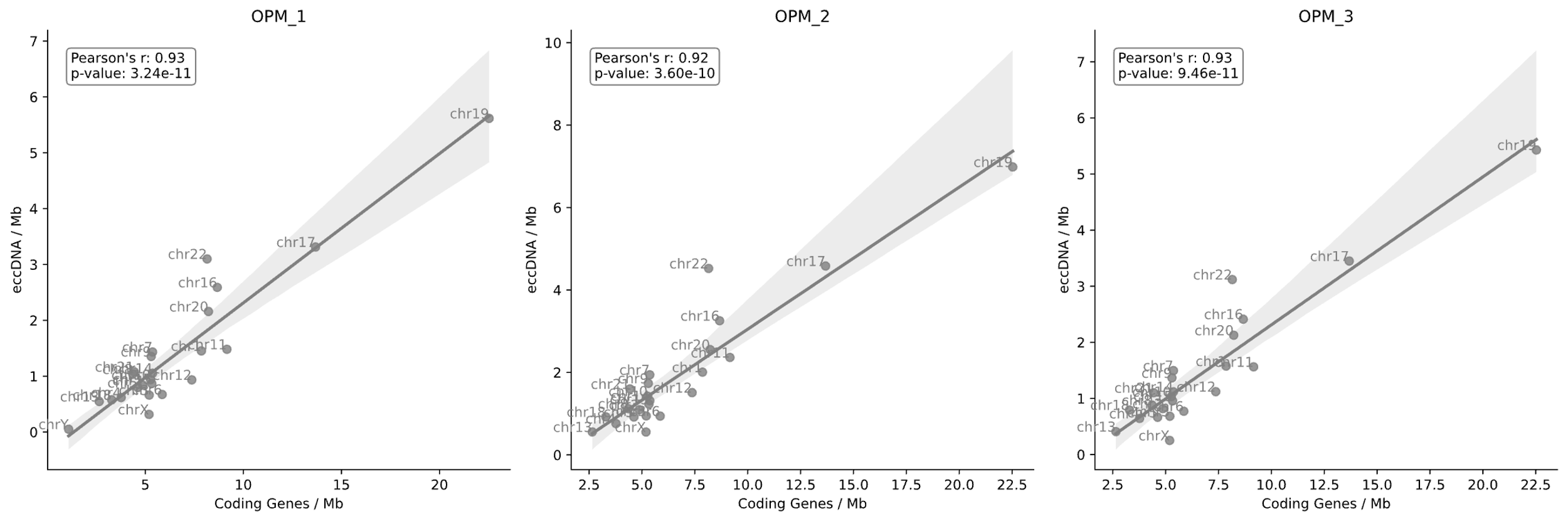

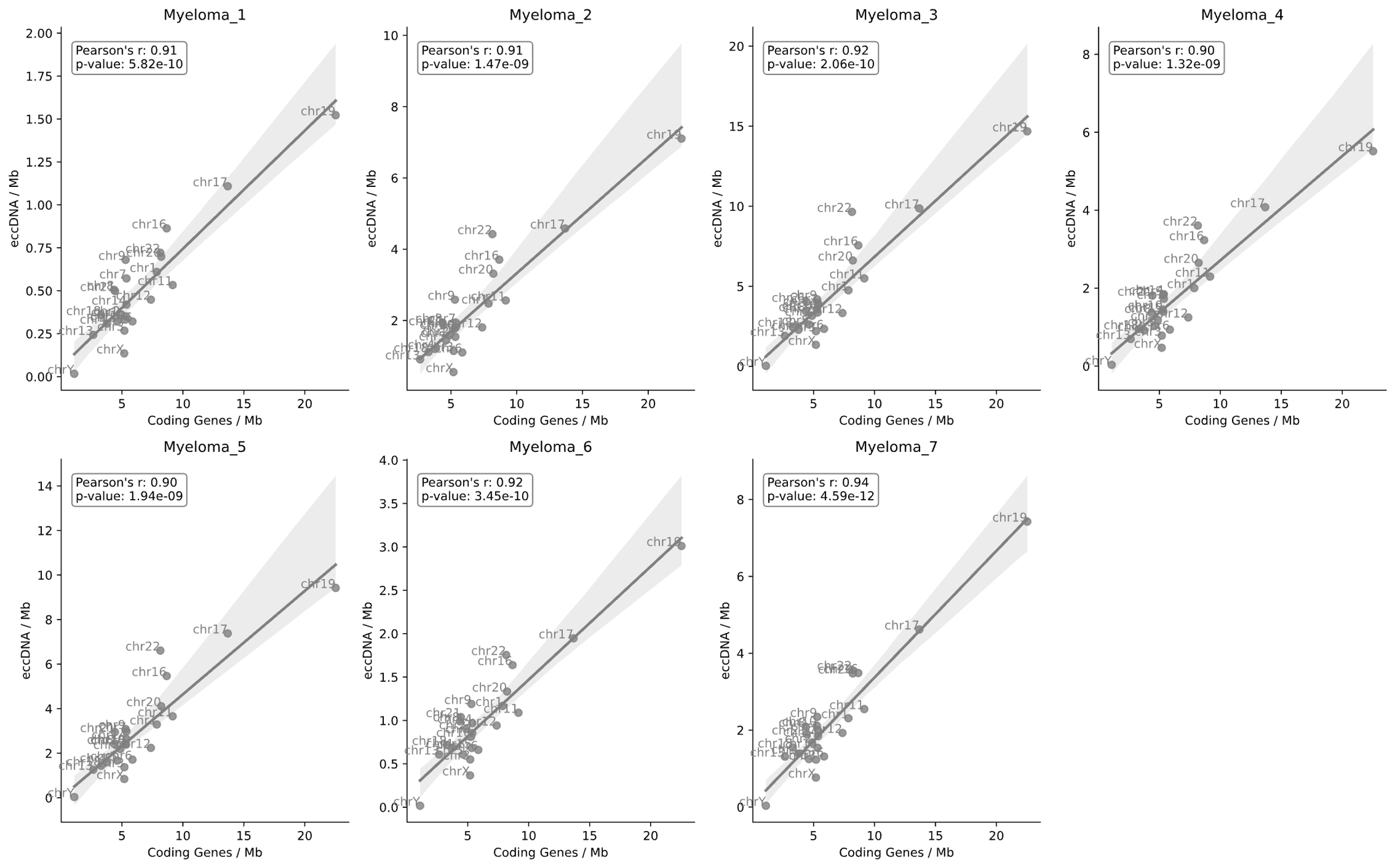

Figure S3

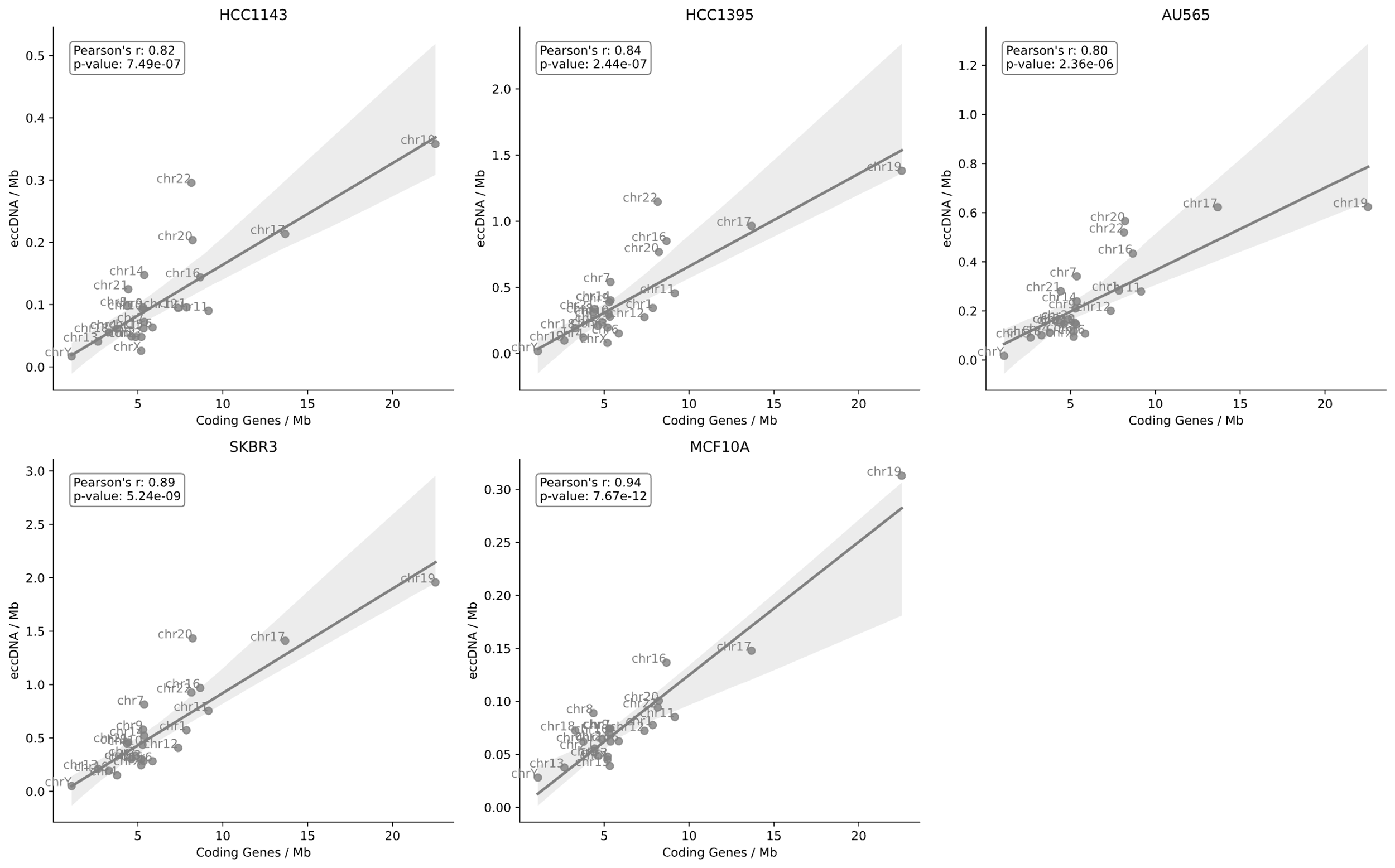

Figure S4

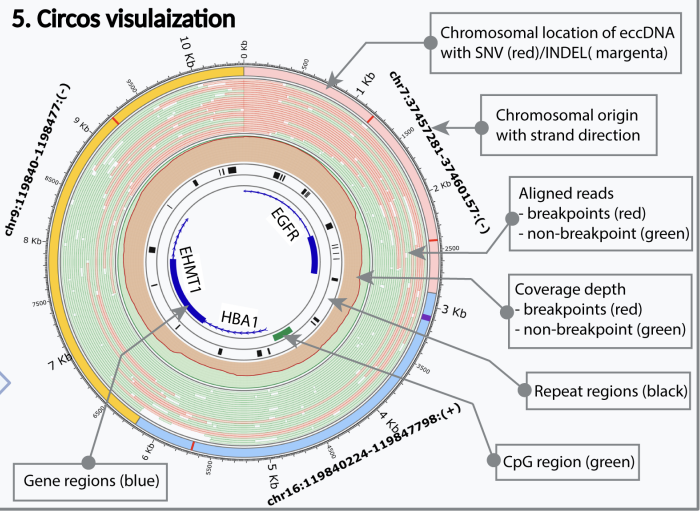

**Figure S5**

Circos lanes annotation from (<https://academic.oup.com/bib/article/23/6/bbac422/6747811> figure 1)

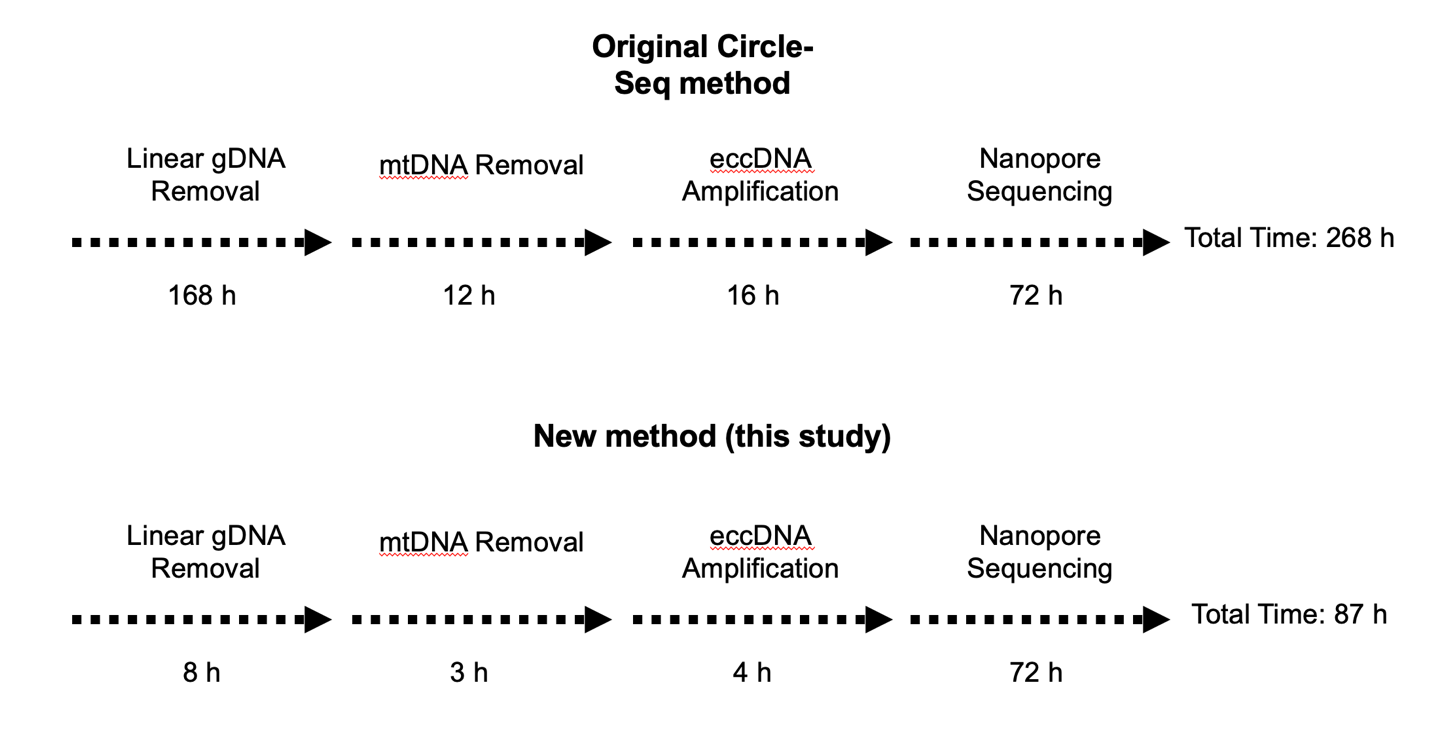

Supplementary workflow

**Table S1**

Amplicon product (µg) of different primer concentrations based on 1 µg of starting gDNA.

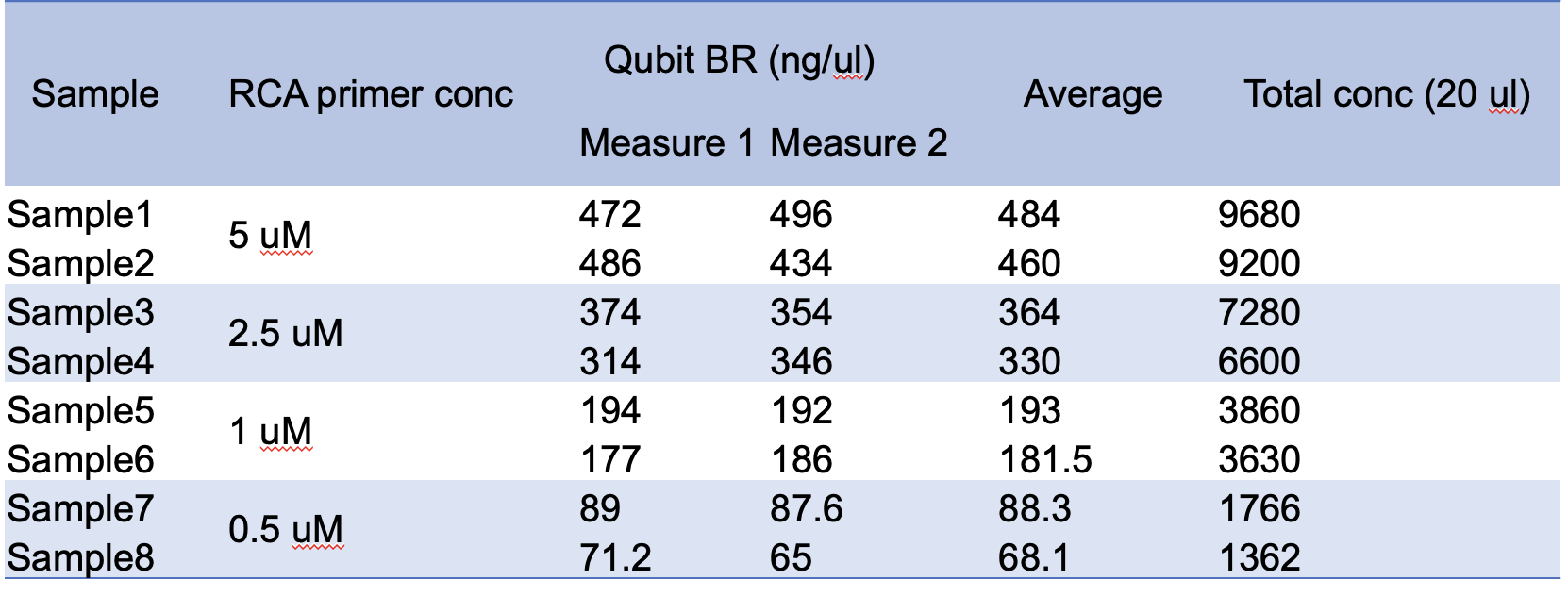

Table S2 Example of eccDNA containing restriction site of either PemI or PacI

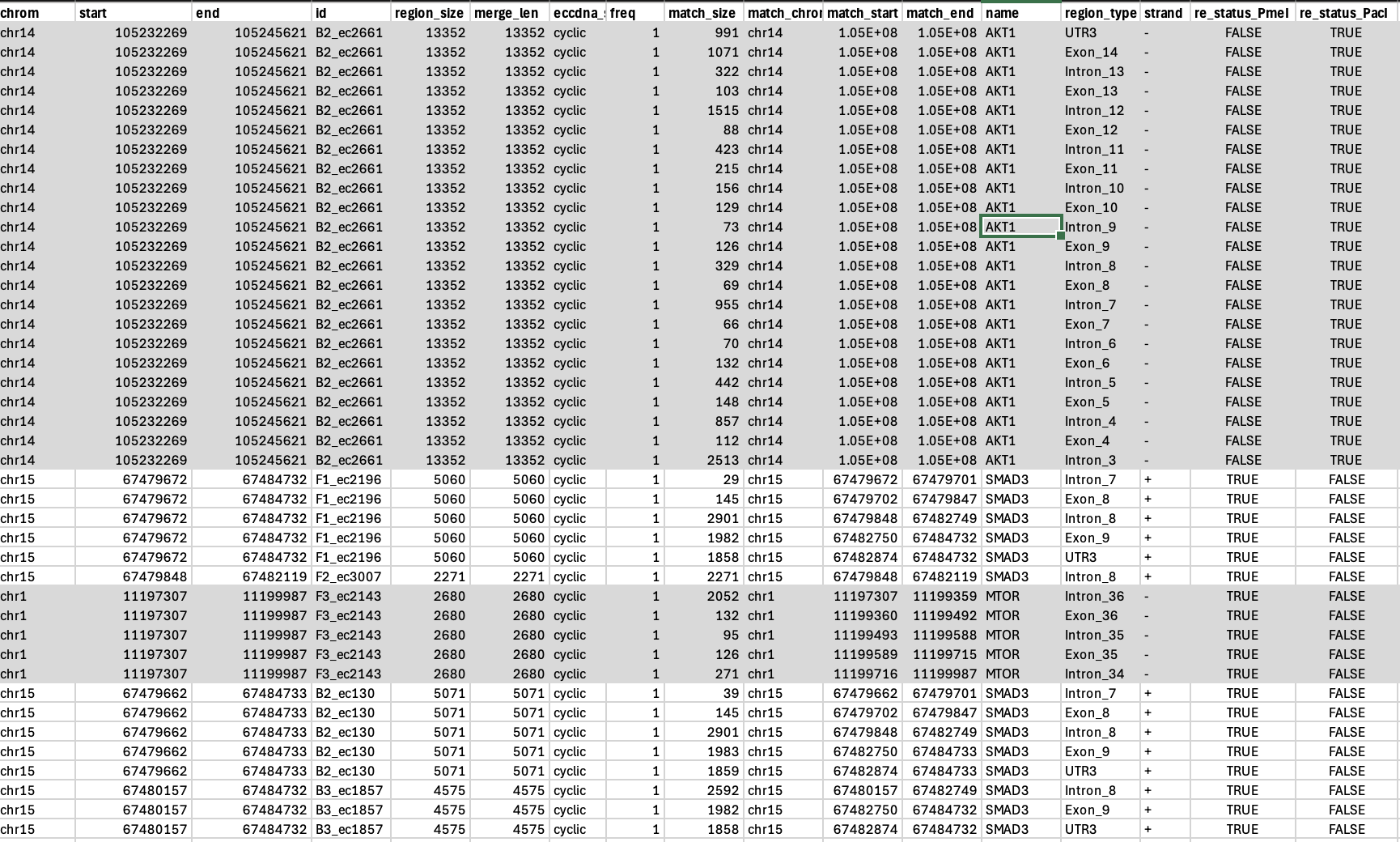
